# Ewing sarcoma breakpoint region 1 prevents transcription-associated genome instability

**DOI:** 10.1101/034215

**Authors:** Nitish Mittal, Christophe Kunz, Foivos Gypas, Shivendra Kishore, Georges Martin, Friedel Wenzel, Erik van Nimwegen, Primo Schär, Mihaela Zavolan

## Abstract

Ewing Sarcoma break point region 1 (EWSR1) is a multi-functional RNA-binding protein that is involved in many cellular processes, from gene expression to RNA processing and transport. Translocations into its locus lead to chimeric proteins with tumorigenic activity, that lack the RNA binding domain. With crosslinking and immunoprecipitation we have found that EWSR1 binds to intronic regions that are present in polyadenylated nuclear RNAs, which include the translocation-prone region of its own locus. Reduced EWSR1 expression leads to gene expression changes that indicate reduced proliferation. By fluorescence in situ hybridization (FISH) with break-apart probes that flanked the translocation-prone region within the EWSR1 locus we found that reduced EWSR1 expression increases the frequency of split signals, indicative of DNA double strand breaks (DSB). The response in phosphorylated histone H2AX and p53-binding protein 1 (53BP1) double-stained foci to the topoisomerase poison camptothecin in cells treated with a control shRNA and with sh-EWSR1 further suggests that EWSR1 functions in the prevention of DNA DSBs. Our data reveal a new function of the EWSR1 member of the FET family and suggest a connection between the RNA-binding activity of EWSR1 and the instability of its own locus that may play a role in malignancy-associated translocations.

## INTRODUCTION

EWSR1 is a member of the FET family of RNA-binding proteins (RBPs), which additionally includes fused in sarcoma (*FUS*) and TATA-box binding protein-associated factor 15 (*TAF15*). These multifunctional proteins have been extensively studied, though mostly for their chimeric variants that are defining features of specific malignancies such as the Ewing’s sarcoma and acute leukemias (reviewed in [1]). FET proteins have a complex domain structure, which includes a glycine/glutamine/serine/threonine-rich transcription activation domain (TAD) at the N terminus, a zinc finger (ZnF) domain at the C terminus and a central RNA recognition motif (RRM). The TAD of EWSR1 further includes an IQ domain that binds calmodulin [2], and a region responsible for the interaction with the SF1 splicing factor [3]. The RRM and ZnF domains are followed by low complexity arginine/glycine-rich (RGG) boxes, structural elements which are found in over a thousand human proteins and mediate interactions with RNAs and proteins [4]. The malignancy-associated chimeric proteins arise through translocations that fuse the locus of a FET protein with a locus encoding a transcription factor. EWSR1 is most commonly translocated to the FLI1 (friend leukemia virus integration 1) transcription factor. The resulting EWSR1-FLI1 fusion protein is a strong transcriptional activator but lacks EWSR1’s RNA-binding domain [5].

A recent study identified many proteins, particularly splicing factors that interact with EWSR1 [1,6]. EWSR1 has also been reported to interact with the hsRPB7 subunit of RNA polymerase II (RNAPII) [7,8] and with another multi-functional pre-mRNA splicing factor, YB-1, that functions in transcription regulation as well as translation [9]. Many of EWSR1’s interactions are sensitive to RNase A treatment, indicating that they are mediated by RNAs [6]. *In vitro*, EWSR1 was reported to bind G- and U-oligomers, through a C-terminal RGG box [10]. *In vivo*, the determination of FET proteins binding specificities lead to seemingly contradictory results. A first photoreactive nucleoside activated (PAR)-CLIP study of FET proteins found that the FUS member of the family binds AU-rich stem-loops (Hoell et al., 2011). However, subsequent studies of FUS reported GU-rich elements [11,12], consistent with *in vitro* determined sequence specificity [10].

Many studies, including those mentioned above, have lead to the view that FET proteins couple various steps in RNA metabolism such as transcription, splicing and transport. The consequences of disrupting this coupling are expected to be broad. However, the phenotypes that were reported in cellular systems with reduced expression of wildtype EWSR1 suggest defects in DNA processing: knockout of EWSR1 in mice led to defects in meiosis and B-lymphocyte maturation [13], whereas the knockdown of EWSR1 in zebrafish led to chromosome segregation defects during mitosis [14]. Whether the EWSR1-dependent RNA processing is involved in DNA replication and cell division is not understood. Rather, what has been reported is a link in the opposite direction, between genotoxic stress and EWSR1 expression and EWSR1-dependent alternative splicing. Induction of DNA damage with camptothecin (CPT) was found to disrupt the interaction between EWSR1 and the spliceosome-associated factor YB-1, leading to many exon skipping events. Among the affected genes was the p53 ubiquitin ligase MDM2, whose increased expression after the removal of camptothecin may lead to the rapid degradation of p53 [15]. Furthermore, irradiation of cells with ultraviolet light was reported to lead to EWSR1 accumulation in the nucleoli and to alternative splicing changes that mirror those induced by depletion of EWSR1 [16]. Some of the affected genes, ABL1, CHEK2, and MAP4K2, are themselves involved in genotoxic stress signaling, which may explain the reduction of viability and proliferation in EWSR1-depleted cells [16]. It remains unclear whether the effects of EWSR1 on the splicing of DNA-damage response genes are mediated by its sequence-specific RNA binding. Very recently, depletion of EWSR1 has also been found to reduce sensitivity to Fas-induced apoptosis [17].

To investigate the relationship between EWSR1’s binding to RNAs and DNA damage, we first re-evaluated the RNA targets of EWSR1 with PAR-CLIP. Consistent with studies of the related protein FUS [11,12], we found that EWSR1 binds broadly along transcription units, particularly in introns, rather than at narrow binding sites. Sequencing of polyadenylated nuclear RNAs revealed a correlation between the density of reads that were captured from intronic regions in EWSR1-CLIP and in nuclear mRNA-seq. In both PAR-CLIP and mRNA-seq samples we observed a very high density of reads originating from the intronic regions of FET family genes where the cancer-associated translocations occur, suggesting a relationship between the RNA binding of EWSR1 and DNA lesions. Further supporting this relationship, we found that shRNA-mediated knock-down of EWSR1 increased the frequency of DNA double-strand breaks (DSBs) in untreated cells but did not affect the level DSBs induced by the topoisomerase-inhibiting drug camptothecin. Strikingly, EWSR1 depletion resulted in instability at the recombination-prone region of the EWSR1 locus. Our data thus suggests that binding of EWSR1 to RNAs acts to prevent transcription-associated genome instability.

### Results

#### EWSR1 binds broadly within pre-mRNAs

The *in vivo* binding specificity of EWSR1 was previously determined in EWSR1-overexpressing HEK293 cells [18]. To determine the targets of the endogenously-encoded EWSR1 protein, we carried out PAR-CLIP (Figure 1A) in HeLa cells. Sequencing of the derived deep sequencing library yielded 78’138’830 reads, which we mapped to the human genome and to transcripts from the release 75 of ENSEMBL with an updated version of the CLIPZ server [19]. Within annotated gene loci, 89% of reads mapped to intronic regions and only 11% to exons. 53.8% of all mutations identified in the reads were T-to-C transitions, as expected from PAR-CLIP [20,21]. To compare with published data sets and to assess the reproducibility of our data we also crosslinked and immunoprecipitated the endogenous EWSR1 in HEK293 cells. The SDS-PAGE gel for this sample showed two bands, one at ~80 kDa and the other at ~120 kDa (Figure 1A), and we used them separately for sample preparation. The summary of these data sets is shown in Supplementary Table 1. To assess the similarity between samples, we computed the distribution of Pearson correlation coefficients between the position-dependent read coverages of individual genes in pairs of samples. To account for the possibility that EWSR1 may interact with the RNA polymerase and thereby bind beyond gene boundaries, we extended the loci by 1000 nucleotides upstream and downstream of the annotated genes. The results are shown in Figure 1B. They indicate, first, that replicate samples are most highly correlated, in both our data set and the data of Hoell et al. [18], as expected. CLIP samples obtained in HEK cells for EWSR1 and for the 3’ end processing factor CFIm, which also binds broadly within transcription units [22] have the next highest correlation, followed very closely by EWSR1-CLIP samples that we prepared in the two different cell types, HeLa and HEK. The correlation between CLIP experiments carried out in HEK293 cells by different labs is also statistically significant (p-value < 2.2e-16 in the Kolmogorov-Smirnov test comparing the Pearson correlation coefficients of positional read coverages in our EWSR1 CLIP and the CLIP of Hoell et al. and the similarly computed Pearson correlation coefficients between the positional coverages in either EWSR1 CLIP data set and randomized data). However, data obtained in different labs show lower correlation compared to replicate data sets obtained in the same laboratory, most likely due to differences in the protocol (see below). The correlation between the read coverage in ELAVL1-CLIP (ELAVL1 is an RBP that interacts with RNAs both in the nucleus and cytoplasm, but is not associated with RNAPII) and EWSR1-CLIP that are performed in the same cell line and same laboratory is at the similar level to the correlation between EWSR1-CLIP performed in the same cell line but in different laboratories.

**Figure 1.**
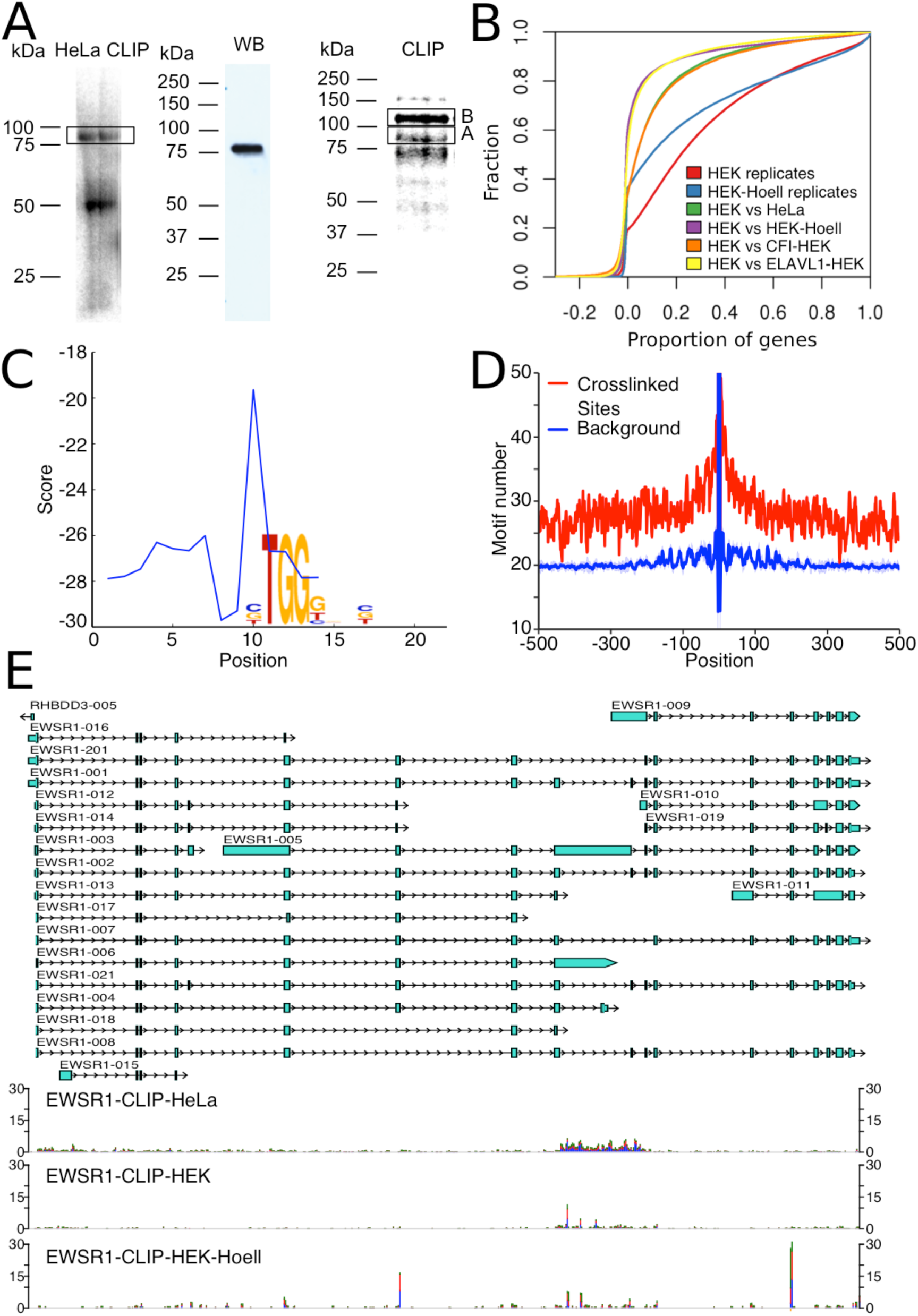
PAR-CLIP-based identification of EWSR1 targets. **A**. SDS PAGE showing separation of the immunoprecipitated EWSR1 protein labeled by UV-crosslinked and radioactive RNA fragments. Left panel and right panels show samples prepared from HeLa and HEK 293 cells, respectively. The middle panel shows Western blot analysis of endogenous EWSR1 protein in HeLa cell extracts. Boxes indicate bands used for sample preparation **B**. Cumulative distributions of Pearson correlation coefficients between the position-dependent read coverages of individual genes in the indicated pairs of samples. Gene loci were extended by 1000 nucleotides upstream and downstream of the annotated genes. **C**. The most significantly enriched motif identified in the 21-nucleotide-long regions centered on the top 5000 most significant crosslinked, as well as its positional preference with respect to the crosslink-diagnostic T-to-C mutation (position 11). **D**. Position-dependent frequency of TGG motifs around the 1’000 highest-confidence EWSR1-crosslinked sites (red) that had a TGG trinucleotide at the crosslinked T/U nucleotide and similar for the same number of random genomic regions centered on TGG motifs (blue). 100 randomized data sets were used to construct the background profile and the light blue lines show the standard deviation computed from these sets. **E**. Screenshot of the EWSR1 locus illustrating annotated transcripts from the ENSEMBL database (top) together with the read density in PAR-CLIP samples prepared from HeLa and HEK293 cells in our laboratory as well as a PAR-CLIP sample from HEK 293 cells that overexpress the EWSR1 protein [18] (bottom, project accession SRA025082, sample accession SRS117978)).

#### EWSR1 has a preference for G-rich elements *in vivo*

To characterize the sequence specificity of EWSR1 we first identified genomic sites with a T-to-C mutation frequency that is most consistent with PAR-CLIP [23] and then applied the Phylogibbs algorithm [24] to the 21-nucleotide regions centered on the 1000 most significant, non-overlapping crosslinked sites. We further refined the motif based on a much larger set, of the top 5000 crosslinked sites, as described in the Methods. Figure 1C shows the most enriched motif as well as its positioning with respect to crosslink-diagnostic T-to-C mutations. The motif is G-rich, consistent with the results of *in vitro* binding studies [10], but different from the stem-loop proposed by Hoell et al. for the over-expressed EWSR1 protein [18]. The reason for this discrepancy is most likely in the PAR-CLIP sample preparation; as has been noted before [21], the protocol used by Hoell et al. [18] appears to lead to the selective loss of G nucleotide-containing sites. Further arguing for this interpretation is the similar discrepancy between the sequence specificity that was reported for the EWSR1 paralog FUS by Hoell et al. and a third group [11]. G-rich nascent RNA can form stable hybrids with the template DNA strand, leading to so-called R-loops. These have been linked to genomic instability due to collisions between various molecular machines that operate on the DNA (see [25] for a review). To determine whether the G-rich motif that we obtained is enriched over broader regions around the crosslinked sites, we used the set of 1’000 highest-confidence crosslinked positions with a TGG trinucleotide (T/U being the crosslinked nucleotide), extracted genomic sequences of 1’001 nt centered on the crosslinked position and computed the position-dependent frequency of occurrence of the TGG motif. We repeated the procedure for 100 sets of the same number (1’000) of random genomic positions where a TGG trinucleotide occurred. As shown in Figure 1D, the density of TGG trinucleotides is 1.5 to 2 fold higher in the crosslinked regions, at least up to 500 nucleotides on either side of the crosslinked position. Thus, EWSR1-bound regions are relatively G-rich, and may have the potential to form R loops.

#### EWSR1 is captured on intronic regions from polyadenylated nuclear mRNAs

We used the set of 20’000 highest-confidence crosslinked positions to identify the EWSR1 targets in which the largest number of crosslinks reside. Strikingly, among these genes are those that themselves encode the FET proteins: FUS ranked 12^th^, with 186 of the top 20’000 crosslinked positions, and EWSR1 ranked 24^th^, with 70 crosslinked positions. The nuclear speckle-associated lincRNAs NEAT1 and MALAT1 had the highest number of high-confidence crosslinked positions, 1938 and 1247, respectively. Supplementary Table 2 contains the annotated top 20’000 sites. That FET proteins crosslink to their own loci was also observed by Lagier-Tourenne et al. [11]. What we additionally find here is that the regions in the FET genes, and in particularly in EWSR1, that are most densely covered by EWSR1 crosslinks are precisely those where translocations occur in malignancies ([26,27], see also http://tinyurl.com/nu9vosk[26]. Figure 1E and Supplementary Figures 1 and 2 show the coverage of these loci in our data, as well as in the data that Hoell et al. obtained for the overexpressed EWSR1 protein in HEK293 cells.

**Figure 2.**
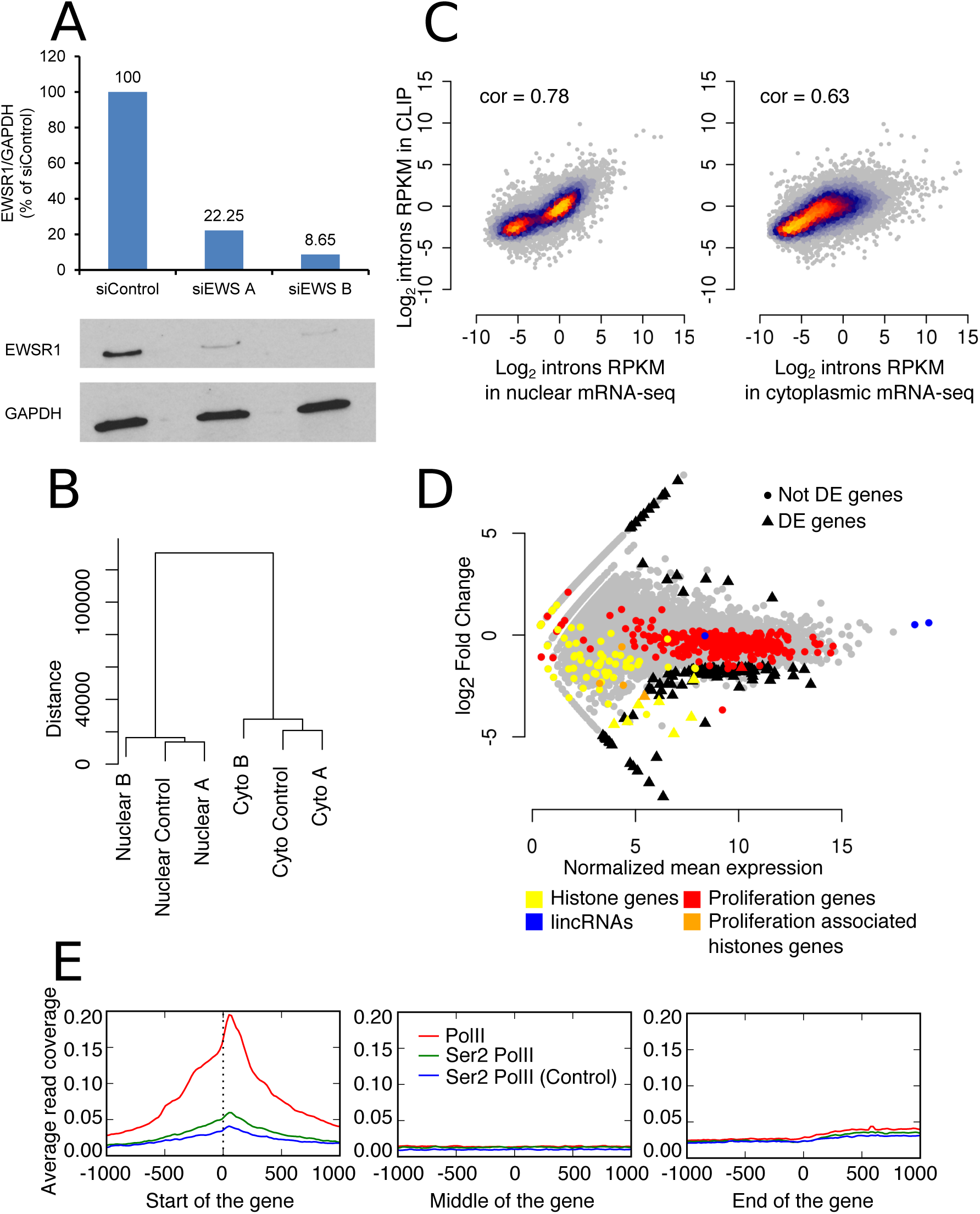
Analysis of mRNA expression upon siRNA-mediated knock-down of EWSR1. **A**. Western blot showing the reduced expression of the EWSR1 protein upon use of two different siRNAs. **B**. Clustering dendrogram of transcript level estimates from nuclear and cytoplasmic fractions of cells treated with a control siRNA or with two distinct siRNAs directed against the EWSR1 transcript. **C**. Scatter plots of the coverage of intronic regions by PAR-CLIP reads and mRNA-seq reads prepared from either the nuclear (left panel) or the cytoplasmic (right panel) fraction. The respective Pearson correlation coefficients are indicated. **D**. MA-plot showing the mean expression (x-axis) and log_2_ fold-change expression of genes in the nuclear fractions of cells treated either with a control siRNA or siRNAs targeting EWSR1. Differentially expressed genes (adjusted p-value < 0.1) are shown with triangle symbols, other genes are shown with circles. Genes whose expression correlates has been associated with the proliferation rate of cells are shown in red and histone genes are shown in yellow. Proliferation-related histones are shown in orange. Abundant nuclear lincRNAs (MALAT1, NEAT1, MEN1) are shown in blue. **E**. Average read coverage of regions around transcription start site, gene middle and transcription ends by reads in various RNA PolII ChIP-seq experiments.

To further investigate the properties of EWSR1-bound intronic regions we sequenced polyadenylated mRNAs from the nuclear and cytoplasmic fractions. Moreover, to explore the consequences of EWSR1 binding to target intronic regions, we sequenced nuclear and cytoplasmic polyadenylated mRNAs from HeLa cells that were either treated with a control siRNA or treated with siRNAs targeting EWSR1. We used two distinct siRNAs, to ensure that the observed changes are not due to off-target effects of the siRNAs. Figure 2A shows that siEWS A reduced EWSR1 expression to 22.25% and siEWS B to 8.65%. Although the samples prepared from the nuclear fraction are much more similar to each other than the samples prepared from the cytoplasmic fraction, the overall gene expression in si-EWSR1-treated cells is not distinguishable from that of control siRNA-treated cells, indicating that EWSR1 does not have a global effect on gene expression levels (Figure 2B). As expected from the localization of the protein and from EWSR1 binding predominantly to intronic regions (Figure 1E), we found that intron coverage in EWSR1-PAR-CLIP was more strongly correlated with the intron coverage in nuclear mRNA-seq compared to intron coverage in cytoplasmic mRNA-seq (Pearson correlation coefficient r = 0.78 vs r = 0.63, p-values < 2.2e-16, Figure 2C). Furthermore, the partial correlation of intron coverage in EWSR1-PAR-CLIP and nuclear mRNA-seq was highly significant (r = 0.58, p-value ~ 0) whereas it was absent between EWSR1-PAR-CLIP and cytoplasmic mRNA-seq. These results suggest that EWSR1 binds to intronic regions that are present in polyadenylated nuclear mRNAs, presumably because they splice at very low rate. As Figure 1E shows, the curated EWSR1 transcripts that are deposited in ENSEMBL show a variety of processing patterns at the intronic region where EWSR1 binds, including what appears to be intron retention. This region, which corresponds at the genomic level to the translocation-prone region in the EWSR1 locus, can either be spliced out, or included in a transcript, but can also be associated with transcription termination or initiation. This processing pattern suggests complex regulatory influences converging on these gene regions.

#### Gene expression changes upon EWSR1 knock-down reflect a role in DNA transactions

Although the siRNA-mediated knock-down of EWSR1 does not have a large effect on gene expression, we asked whether any genes change consistently in the EWSR1 knock-downs with respect to the control siRNA transfection. As shown in Figure 2D, siRNA-mediated knock-down of EWSR1 induces consistent down-regulation of 108 genes, whereas only 19 genes are up-regulated. Gene Ontology [28] analysis of the genes that are differentially expressed upon EWSR1 knockdown reveals terms such as nucleosome, DNA packaging complex and protein-DNA complex (Table 1), all related to chromatin structure. Surprisingly, many histone-encoding mRNAs were captured in the poly(A)+ samples (Figure 2D, yellow and orange symbols), in spite of the fact that histone mRNAs are not generally polyadenylated. In addition, the abundance of histone mRNAs was lower in the nuclei from si-EWSR1-treated cells compared to control siRNA-treated cells. We therefore wondered whether this observation may be due to differences in the isolation of nuclear RNA in these experiments. Abundant nuclear non-coding RNAs such as MALAT1, NEAT1 and MEN1 had a similar expression in the control siRNA and si-EWSR1-treated samples (Figure 2D, blue symbols), indicating that differences in the efficiency of isolating nuclear RNA do not explain the depletion of histone mRNAs in si-EWSR1-treated samples. An alternative explanation could be that the depletion of EWSR1 alters the proliferation rate of the cells and thereby the expression of histone genes. We used a set of genes whose expression is indicative of the proliferation rate [29] to evaluate the proliferation status of cells that were treated with various siRNAs. Some of these genes are indeed histones (Figure 2D, orange symbols). Treatment of cells with an EWSR1 specific siRNA reduced expression of proliferation-related genes (Figure 2D, red and orange symbols), indicating that changes in histone gene expression reflect an effect on cell cycle progression.

**Table 1.** Gene Ontology terms that are enriched in genes that undergo differential expression upon EWSR1 knockdown. Genes identified as differentially expressed based on the nuclear mRNA-samples were used in this analysis.

| Gene Ontology category | p-value | Fold enrichment | DE genes in category |
| --- | --- | --- | --- |
| GO:0000786<br>nucleosome | 2.896535e-09 | 34.63 | HIST2H2BE, H1FX, HIST1H2AG,<br>HIST1H2BK, HIST1H1C, HIST1H2AC,<br>HIST1H2BN |
| GO:0044815<br>DNA packaging complex | 1.539988e-08 | 30.92 | HIST2H2BE, H1FX, HIST1H2AG,<br>HIST1H2BK, HIST1H1C, HIST1H2AC,<br>HIST1H2BN |
| GO:0032993<br>protein-DNA complex | 3.994154e-06 | 15.46 | HIST2H2BE, H1FX, HIST1H2AG,<br>HIST1H2BK, HIST1H1C, HIST1H2AC,<br>HIST1H2BN |
| GO:0007176<br>regulation of epidermal growth<br>factor-activated receptor<br>activity | 1.257601e-05 | 52.08 | ERRFI1, SOCS5, EPGN, SOCS4 |
| GO:0061097<br>regulation of protein tyrosine<br>kinase activity | 1.795158e-05 | 26.89 | ERRFI1, SOCS5, EPGN, SOCS4 |
| GO:0007175<br>negative regulation of epidermal<br>growth factor-activated receptor<br>activity | 2.960969e-05 | 92.76 | ERRFI1, SOCS5, SOCS4 |

Although only very few EWSR1 target genes are differentially expressed (Supplementary Table 3), the chi-square test indicates a weak dependency between EWSR1 binding and the response of the targets upon si-EWSR1-mediated knock-down (Supplementary Table 4, chi-square p-value = 0.003336). The annotation of the top 20’000 EWSR1 crosslinking positions revealed that many of these correspond to non-coding RNAs (Supplementary Table 2), including the highly abundant lincRNAs NEAT1 and MALAT1 that localize to nuclear speckles, sites of pre-mRNA splicing factor enrichment [30]. Also EWSR1 appears to bind strongly to these lincRNAs, their expression is not affected upon si-EWSR1-mediated knock-down. Small non-coding RNAs such as miRNAs and snoRNAs that are processed from the introns of pre-mRNAs are also represented among the high-confidence targets of EWSR1. These associations may explain previously reported changes in pre-mRNA splicing when EWSR1’s expression is perturbed [15,16].

#### EWSR1 knock-down increases the occupancy of gene start regions by Ser2-phosphorylated RNA polymerase II

A previous study of the FET family member FUS found that this protein regulates the Ser2-phosphorylation of RNAPII’s carboxyterminal domain [31]. To determine whether EWSR1 shares this function, we determined the coverage of transcription units by the elongating form of RNA PolII, whose carboxyterminal domain is phosphorylated at serine 2, in control siRNA or si-EWSR1-treated cells. The coverage of transcription units by the RNA PolII has the expected profile, with peaks around the transcription start sites and in the region immediately downstream of gene ends (Figure 2E). Moreover, Ser2-phosphorylated RNA PolII density around the start of gene loci shows a slight increase in the si-EWSR1-treated cells compared to control siRNA-treated cells. This suggests that EWSR1 promotes elongation by RNA PolII. A summary of the EWSR1 locus coverage by RNA PolII ChIP reads is shown in Supplementary Figure 3.

**Figure 3.**
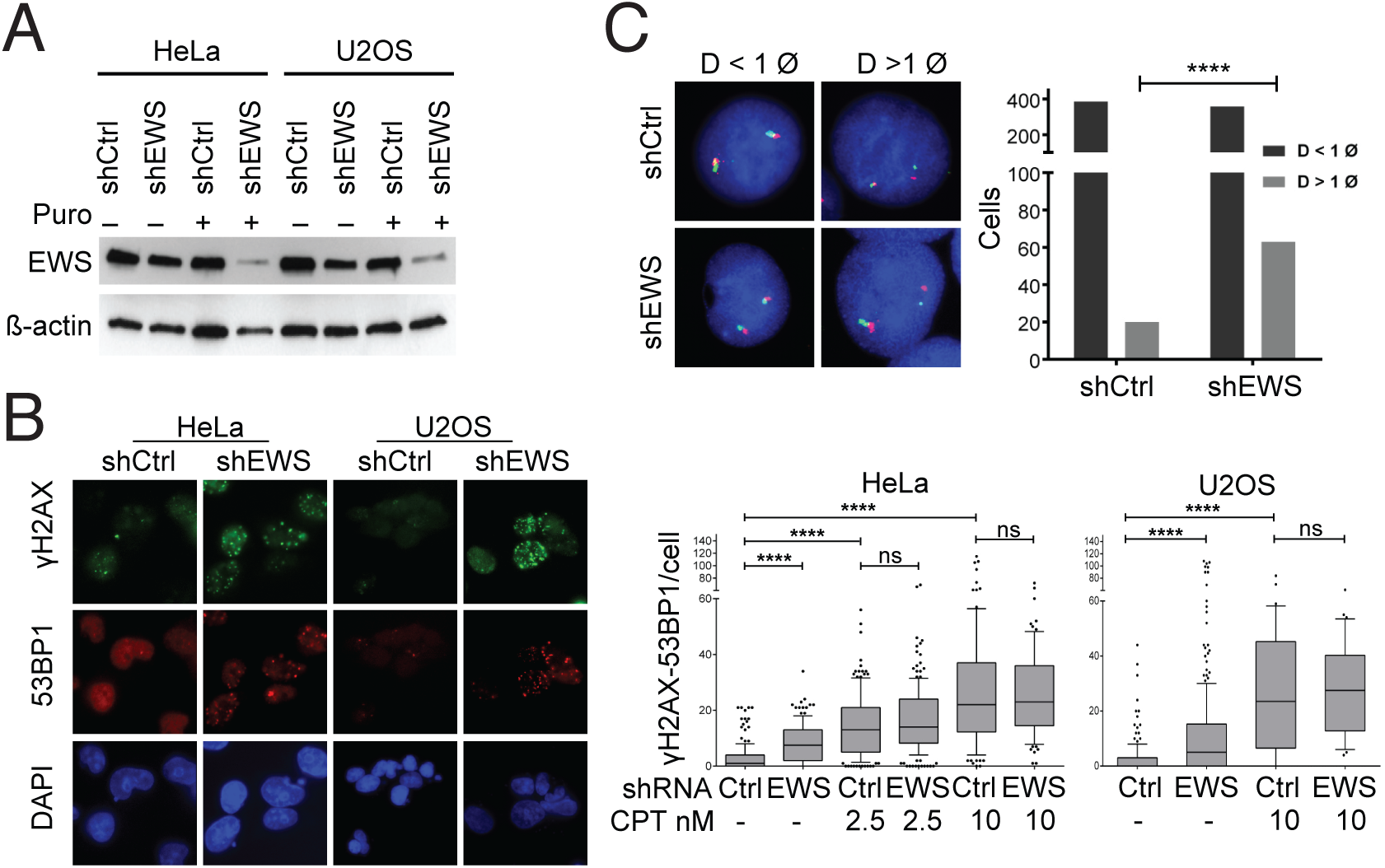
EWSR1 prevents genomic instability. **A**. Western blot showing EWS and β-actin protein levels in HeLa and U2OS cells treated with EWS or control shRNAs four days after transfection with and without mild puromycin (Puro) selection. **B**. Representative immunofluorescence images and quantification of spontaneously formed nuclear γH2AX/53BP1 co-foci in HeLa and U2OS cells enriched for expression of EWSR1 or control shRNA. CPT indicates additional camptothecin treatment. DNA was stained with DAPI. Box plots summarize the number of γH2AX/53BP1 co-foci per cell. 10–90 percentiles values are shown as whiskers. ****: p < 0.0001 in the Kruskal-Wallis test; ns: not significant. **C**. Representative FISH images and the quantification of EWS break-apart probe signals showing intact (D < 1 ø) or split (D > 1 ø) patterns in the *EWS* locus in HeLa cells expressing EWS or control shRNA. ****: p < 0.0001 in Fisher’s exact test.

Altogether, the results described in the previous sections lead us to hypothesize that the EWSR1 may promote transcription, particularly at genes with complex structure. If this were the case, we would expect EWSR1 target regions to be susceptible to transcription-associated stress and DNA breaks.

#### EWSR1 acts in the prevention of genome instability

To determine whether EWSR1 depletion increases the frequency of DNA DSBs, we determined the frequency of phosphorylated H2AX and 53BP1 co-staining nuclear foci following transient short hairpin RNA (shRNA) mediated knockdown of EWSR1. These proteins are recruited at sites of non-homologous end joining (NHEJ) repair of DSBs [32]. Relative to a control shRNA, the EWSR1-specific shRNA markedly reduced EWSR1’s protein levels in both HeLa and osteosarcoma (U2OS) cells successfully transfected with the shRNA constructs (Figure 3A). The reduction in expression was also accompanied by a significant increase in the frequency of γH2AX/53BP1 double-stained nuclear foci in both HeLa and U2OS cells (Figure 3B; Kruskal-Wallis test p-value<0.0001). That is, the median number of double-stained foci increases from 1 or 0 to 7.5 or 5, respectively, in HeLa and U2OS cells. Treatment of cells with a high dose of camptothecin (CPT), a DNA damaging agent that acts as a topoisomerase 1 poison to cause transcriptional arrest and replication fork collapse [33] also led to an increase in the median number of double-stained foci to 22 and 23.5 in HeLa and U2OS cells, respectively, while a sub-lethal dose of CPT had a moderate effect, comparable to the EWSR1 knock-down (median number of double-stained foci of 13 in HeLa cells). The shRNA mediated knock-down of EWSR1 did not further increase the number of double-stained foci, neither in cells treated with a high (HeLa: 22 vs 23; U2O2: 23.5 vs 27.5) nor with a sublethal dose of CPT (HeLa 13 vs 14) (Figure 3B; Kruskal-Wallis test p-value=non-significant (ns)). Thus, the knock-down of EWSR1 significantly increases the steady state levels of γH2AX/53BP1 double-stained foci in HeLa and U2OS cells, but it does not further increase the frequency of CPT-induced foci. These results suggest that EWSR1 may function in the prevention of DNA DSBs.

To establish that EWSR1 indeed prevents genome instability at regions where it binds its RNA targets, we carried out FISH with EWSR1 break-apart probes that were designed to flank the region between intron 7 and intron 10 of the EWSR1 locus which is the translocation hotspot. We then measured the number of split signals, indicative of DSBs, in HeLa cells that were either treated with a control or with an EWSR1-specific shRNA. We found that EWSR1 depletion lead to a significant, 3-fold increase in the frequency of cells with one or more split signals (Figure 3C; Fisher’s exact test p-value < 0.0001). That is, compared to 20 split signals in control shRNA-treated cells, we observed approximately 60 split signals in EWSR1 shRNA-treated cells. This result suggests that depletion of EWSR1 creates an environment that is permissive for chromosomal translocations at its own genomic locus, where EWSR1 strongly associates with RNA.

## DISCUSSION

FET proteins are complex, multi-functional proteins that have been implicated in a broad range of processes, including transcription, splicing, mRNA translation as well as DNA damage signaling. To better understand how binding of EWSR1 to RNAs may affect DNA damage prevention or repair we have first determined the *in vivo* targets of EWSR1 and established that EWSR1 crosslinking is strongly enriched in the intronic regions of FET protein transcription where the DNA is targeted for translocation in malignancies such as the Ewing’s sarcoma. We then studied the behavior of the EWSR1 locus upon shRNA-mediated knock-down of EWSR1 and found an increased frequency of DNA breaks, indicating a role of EWSR1 in the prevention of transcription-associated DNA breaks.

The binding specificity of EWSR1 was studied initially *in vitro*, where EWSR1 was reported to bind G- and U-runs [10]. Recently, a number of studies employed ultraviolet light-mediated crosslinking and immunoprecipitation to identify FET protein targets. Hoell et al. studied all wildtype forms of FET family members as well as a few variants that are associated with amyotrophic lateral sclerosis [18]. Rather than the G/U-rich motif that emerged from *in vitro* studies, this study proposed that FET proteins bind stem loop structures. Lagier-Tourenne et al. [11] and Rogelj et al. [12] studied only the FUS member of the family and reported that FUS binds G/U-rich motifs, consistent with the *in vitro* data. In our study, we re-assessed EWSR1’s binding specificity with PAR-CLIP and retrieved a G/U-rich motif as well. The discrepancies in the *in vivo* data are most likely due to differences in the protocols that were used in CLIP experiments. In particular, extensive T1 RNase digestion (a ribonuclease that cleaves after guanine residues) could lead to the strong depletion of G-containing binding sites [21]. In contrast, our results, obtained with mild digestion with RNase I (a ribonuclease that does not have strong nucleotide specificity), are consistent with both the previous *in vitro* data obtained for EWSR1 as well as with other CLIP studies of the related FUS protein. Thus, binding of G/U-rich elements *in vivo* is most likely a conserved feature of FET family proteins.

EWSR1 was previously linked to alternative splicing [15,16,34]. The mechanisms underlying splicing changes are likely multiple. EWSR1 has protein-protein interactions with many splicing regulators [1] and interestingly, here we found that many RNAs that are bound by EWSR1 encode splicing regulators. A recent study found that the FUS protein regulates the phosphorylation of RNAPII at serine 2 and thereby the rate of RNAPII elongation [31]. In turn the rate of RNA PolII elongation has been shown to modulate splicing (reviewed in [35]). Here we have observed a small difference between the density of Ser2-phosphorylated RNA PolII ChIP reads in EWSR1 siRNA-treated cells compared to control siRNA-treated cells. Although the mechanisms underlying the changes in splicing upon EWSR1 depletion need further studies, it is possible that EWSR1 contributes to the recognition of specific intronically-encoded RNAs such as miRNAs, snoRNAs and others, and influences the rates of various competing RNA processing steps, including transcription. In support of this hypothesis comes the general down-regulation of mRNAs, whether or not targets of EWSR1, upon EWSR1 knock-down. Interestingly, histones are among the genes that are significantly down-regulated upon si-EWSR1 treatment, an observation that was previously made for the FUS paralog of EWSR1 [36]. FUS appears to interact with the U7 snRNA and to regulate both transcription and the 3’ end processing of histone genes. The FUS interactions with histone gene expression machinery occur during specific phases of the cell cycle. Our EWSR1-CLIP in HeLa cells captured as well some reads from the U7 snRNA. However, the involvement of EWSR1 in the regulation of histone gene processing could be addressed in depth in future studies.

The EWSR1 protein is best known for its fusion forms that are characteristic for Ewing sarcomas and are due to chromosomal translocations. The most common translocation, observed in 85% of Ewing sarcomas, involves the *FLI1* locus. It leads to the production of the EWS/FLI1 fusion protein that lacks the RNA binding domain of EWSR1 [10], and has strong transcriptional activity. Surprisingly, the observed increase of DSB signaling as well as our FISH analysis suggests that reduction of EWSR1 levels could lead to DNA breaks in its own translocation-prone locus, and more generally, to DSBs. Whether an initial reduction of EWSR1 level is part of the pathogenic process of Ewing sarcoma is an intriguing hypothesis for future studies.

### Methods

#### PAR-CLIP experiment and analysis

We mapped binding of EWSR1 to RNA *in vivo* by a modified photoreactive nucleotide-based CLIP (PAR-CLIP) method ([37], adapted by Martin et al., [22]). To this end, Hela cells were grown in 15 cm dishes in the presence of 4-thiouridine and RNA-protein crosslinks were generated by exposure to 365 nM UV-light. EWSR-containing complexes were precipitated with a mouse monoclonal antibody against EWSR1 (C-9, sc-48404; Santa Cruz Biotech), and crosslinked RNA fragments were extracted from the isolated complexes, ligated to adapters and amplified by RT-PCR. Sequencing was performed on an Illumina HiSeq 2500 platform for 50 cycles and sequence reads were pre-processed and mapped to the human genome using CLIPZ[19].

We then used the procedure described in [23] to identify genomic positions for which the T-to-C conversion frequency determined from the sequenced reads is most consistent with what is expected from a crosslinking site. We sorted the sites based on the posterior probability of the site having a crosslinking-consistent mutation pattern, extracted the top 20000 sites and then clustered those that were within 10 nucleotides of each other. From each of the 11’893 resulting clusters we extracted a representative site, which was the one with the highest posterior probability. We extended these sites by 10 nucleotides on each side and ran the PhyloGibbs motif finding software [38] on the top 5000 sites to identify an initial motif of length 8 (using a 0^th^-order background model, asking for 4000 sites among the 5000 sequences, and restricting sites to occur only on the positive strand, as expected for an RNA binding protein). We then used the MotEvo software [39] to refine this motif on the full set of 11’893 target sequences (using a prior that corresponds to expecting one site per sequence on average, and again restricting sites to occur only on the positive strand).

We then used the 1’000 highest-confidence crosslinked sites that had a TGG trinucleotide at the crosslinked T/U nucleotide and counted the number of occurrences of the trinucleotide in an extended window of 500 bases left and right of the crosslinked nucleotide. The same procedure was repeated for random TGG trinucleotides in the genome. Their sequences were extracted from the genome with bedtools [40]. Profiles for the crosslinked sites (red) and the background (blue) were generated using a sliding window of five nucleotides with the aid of the seaborn library [41]. For the background, we used 100 randomized sets of sequences and the light blue lines show the standard deviation of motif counts in the 100 randomizations.

#### mRNA-seq experiment and analysis

Cytoplasmic and nuclear fraction of HeLa cells transfected with control siRNA and two different si-EWS RNAs were obtained according to Suzuki K et al. [42]. Poly(A)+ RNA was isolated from cytoplasmic and nuclear fractions with Dynabeads^®^ mRNA DIRECT^™^ Kit (61011, life technologies) according to the manufacturer’s protocol. After isolation of mRNA, sequencing libraries were prepared using the “Directional mRNA-seq sample preparation” protocol from Illumina with minor modifications. In brief, isolated mRNA was chemically fragmented by incubating the mRNA solution with twice the volume of alkaline hydrolysis buffer (50 mM Sodium Carbonate [NaHCO_3_/Na_2_CO_3_] pH 9.2, 1 mM EDTA) at 95 °C for 5 minutes to get fragments of approximately 200–350 bases. Fragmented mRNA was immediately purified with RNeasy MinElute Cleanup Kit (74204, Qiagen) to stop the reaction and to remove small RNA fragments (<100 bases). Purified fragmented mRNA was treated with thermo-sensitive alkaline phosphatase FastAP (EF0651, Fermentas) at 37 °C for 30 minutes and then at 75 °C for 5 minutes to inactivate FastAP. Fragmented mRNA was further incubated with ATP and T4 polynucleotide kinase (EK0032, Fermentas) at 37 °C for an hour and subsequently purified. Ligation of RNA 3’ adapter (RA3, part # 15013207, Illumina) was done using T4 RNA Ligase 2, truncated K227Q (M0351L, New England Biolabs Inc) according to the Illumina protocol. The ligation step was followed by RNA purification as mentioned above to remove un-ligated 3’ adapters. The RNA 5’ Adapter (RA5, part # 15013205, Illumina) was ligated using the T4 RNA ligase (EL0021, Fermentas) and then the RNA was purified to remove un-ligated 5’ adaptors. cDNA was synthesized using an RNA RT Primer (RTP, part # 15013981, Illumina) with SuperScript III (18080044, Invitrogen) as per Illumina protocol. Libraries were amplified for 14 cycles of PCR using forward PCR primer (RNA PCR Primer (RP1), part # 15005505 Illumina) and reverse PCR primer (Illumina PCR Primer, Index). Different indexed reverse PCR primers were used for library preparation from different samples to facilitate multiplexing. Libraries were sequenced for 50 cycles on an Illumina HiSeq 2500 instrument.

We trimmed adaptor sequences from the mRNA-seq reads generated from control siRNA and si-EWSR1-treated samples with cutadapt (v1.4.1) [43] with parameters -- adapter TGGAATTCTCGGGTGCCAAGG --error-rate 0.1 --minimim-length 15 -- overlap 1. To obtain transcript and gene expression level estimates we used Sailfish (v0.8.0) [44], a method that we have tested before and found to be fast and accurate [45]. We used ENSEMBL (version 75) transcript sequences including both coding and non-coding RNAs as the transcript set to which reads were mapped and the variational bayesian expectation maximization algorithm (option --useVBOpt) from Sailfish.

Clustering of transcript level estimates (TPM) was performed with R (version 3.2.2) with Euclidean distance between gene expression vectors and the ward.D function for clustering. For the differential expression analysis we used the DESeq package [46] with the rounded estimated number of reads for each gene as input (column NumReads from the Sailfish output file genes.sf). We used a pseudocount of one for read counts associated with a gene. We also used the CLIPZ server [19] to map the reads to the reference genome (hg19) and compare the intronic read coverage (all introns of the longest pre-mRNA) in nuclear and cytoplasmic mRNA-seq samples with the intronic coverage in EWSR1-CLIP.

We used a custom script to select the longest transcript per gene (based on the total intron length) from the ENSEMBL gtf file. We then used bedtools [40] to extract reads whose loci overlapped with the introns of these transcripts (bedtools intersect -f 1 -s -abam <alignment file> -b <annotation>). The alignment files were indexed with samtools [47] and the intronic coverage was estimated with the following command: bedtools multicov -D -s -bams <alignment files> -bed <annotation>.

#### RNA PolII ChIP-seq experiment and analysis

The ChIP Protocol was adapted from Blecher-Gonen et al. [48]. The solutions and steps that differed from the reference are described below. 10^8^ HeLa cells transfected with either control siRNA or with si-EWSR1 were crosslinked in fixing buffer (50 mM HEPES pH 7.5, 0.2 mM EDTA pH 8.8, 0.1 mM EGTA pH 8.8, 100 mM NaCl, 1% formaldehyde) for 10 min with continuous rocking at room temperature, and then quenched with 125 mM glycine for 5 min. Cells were washed thrice with cold PBS and collected by scrapping. Nuclei were isolated and lysed to obtain crosslinked chromatin. Simultaneously, antibodies were coupled with Protein G magnetic beads (88848, Pierce^™^) by incubating 100 μl of Protein G beads with 10 μg of Pol II Antibody (N-20) X (sc-899 X, Santa Cruz biotechnologies) or anti-RNA polymerase II CTD repeat YSPTSPS (phospho S2) antibody (ab5095, Abcam plc), for minimum 1hr at room temperature, with continuous rotation. Further, chromatin was sonicated in cold conditions to reduce the heating effect with a probe sonicator for 6 cycles of 30 sec on and 1min 15 sec off at 60 watt to obtain DNA fragments of 100–500 bp. Fragmented chromatin was centrifuged at 20,000g for 10 min at 4 °C to remove the nuclear debris. 3% chromatin was kept as input from each sample and an equal amount (around 750–1000 μg) of chromatin was incubated with magnetic beads-coupled antibody at 4 °C overnight with continuous rotation. Immuno-complexes were washed with 1 ml of wash buffers as described in the reference. Washed immuno-complexes along with input samples were further treated with RNase and then with Proteinase K followed by overnight reverse crosslinking at 65 °C with continuous shaking at 1400 rpm in a thermoblock with heating lid. DNA was purified using Ampure beads as detailed in the reference.

The sequencing library were prepared according to the instruction manual of NEBNext^®^ ChIP-Seq Library Prep Reagent Set for Illumina. In brief, end repair of input and ChIPed DNA was done by incubating with T4 DNA Polymerase, Klenow fragment and T4 PNK enzyme at 20 °C for 30 minutes. The reaction was purified using Ampure beads according to the instruction manual. An A overhang at the 3’ end was produced by treating end repaired DNA with dATP and Klenow Fragment (3′→5′ exo^−^) at 37 °C for 20 min, after which the DNA was purified. Double stranded DNA adaptors were ligated DNA with dA overhang with the T4 DNA ligase at 37 °C for 30 min. DNA was purified and size-selected as described in the instruction manual. Size-selected DNA was PCR-amplified for 16 cycles using NEBNext High-Fidelity 2X PCR Master Mix with Illumina universal forward primer and indexed reverse primer that facilitate multiplexing of samples for sequencing. Amplified DNA was finally purified and sequenced on an Illumina Hiseq2500 instrument.

Reads were again mapped with the pipeline used in the CLIPZ server, and alignment files (in BAM format) were extracted to generate profiles around the beginning, middle and end of gene loci. We used only data from consensus chromosomes (no haplotypes, starting with H in the ENSEMBL gtf file), and kept only genes whose loci were longer than 4000 bases, to ensure that the 2000 nts-long regions whose coverage we wanted to estimate (from −1000 to +1000 around transcription start, gene middle, and transcription end) do not overlap. We then generated coverage profiles in these regions by Ser2-phosphorylated Poll in the control siRNA and si-EWSR1-treated samples. For reference, we added the profile of coverage by RNA PolII reads. In all cases we subtracted the “background” coverage profile, by reads obtained when sequencing input chromatin. Profiles were generated with the metaseq package [49]. In all cases, read count was normalized by library size.

#### Immunofluorescence

EWS shRNA constructs used were originally obtained from Origene (catalogue number TG313142, sequence shRNA-EWS_1 - GI352561: GAGCAGTTACTCTCAGCAGAACACCTATG) and contained the shEWS sequence cloned in a retroviral GFP vector. shEWS cassette was subsequently sub cloned into a retroviral pRS vector (Origene) without GFP using EcorI/HindIII site to eliminate interference of any GFP fluorescence. HeLa and U2OS cells grown on coverslips in 6-well plates were then transfected with EWS shRNA construct or respective control (sequence TR30013 GCACTACCAGAGCTAACTCAGATAGTACT) in pRS vector containing a puromycin resistance cassette using the jetPEI transfection reagent (Polyplus-transfection SA, France) according to the manufacturers protocol. Medium was exchanged 12 hours after transfection. Successfully transfected cells were then enriched by the addition of 1 g/ml puromycin to the culture medium. Additionally, for continuous CPT treatments, CPT was added to a final concentration of 10 nM. Four days after transfection, the cells were washed twice with PBS and fixed for 30 min in ice-cold methanol at 4°C. All following steps were performed at room temperature. After washing twice with PBS, cells were blocked with blocking buffer (5% FCS in PBS) for 30 min. Antibodies against H2AX (clone JBW301; upstate, USA) and 53BP1 (H-300; Santa Cruz Biotechnology, USA) were hybridized in blocking buffer at dilutions of 1:1000 and 1:500, respectively. After one hour cells were washed three times with PBS for 5 min, and secondary antibodies (A11017 and A11012; Invitrogen, USA) hybridized in blocking buffer for 30 min. After washing 3 times with PBS for 5 min, coverslips were mounted using vectashield, containing DAPI (1 g/ml). H2AX and 53BP1 were evaluated and counted with a Leica DMI 4000 microscope. Statistical analysis of triplicate experiments was done using the Prism5 software (GraphPad, USA).HeLa and U2OS cells transfected with control or EWS shRNA constructs on glass coverslips were enriched by puromycin selection, fixed in ice-cold methanol, stained with antibodies and imaged with a Leica DMI 4000 microscope.

#### Fluorescent in situ hybridization (FISH)

HeLa cells were grown and transfected with control or EWS shRNA constructs as mentioned above. Four days after transfection coverslips were washed twice with PBS and incubated in 2x SSC (pH 7.0) for 30 minutes at 37°C. Samples were then dehydrated by a series of two minutes incubations in 70%, 85% and 100% ice-cold ethanol and subsequently air dried. To avoid over-denaturation in the subsequent steps, samples were “aged” overnight at 4°C. Hybridization of the Poseidon Repeat Free EWSR1 (22q12) Break probe (KBI-10708; KREATECH Diagnostics, Netherlands) and subsequent steps were performed according to the manufacturer’s instructions. Briefly, coverslips were covered with 10 μl of the probe, sealed with Fixogum, denatured at 80°C for 5 minutes and hybridized overnight at 37°C in a humidified chamber. Post-hybridization washes included one time 2xSSC/0.1% Igepal for 2 minutes at room temperature, one time 0.4x SSC/0.3% Igepal for 2 minutes at 72°C and one time 2xSSC/0.1% Igepal for 2 minutes at room temperature, the latter two steps being performed without agitation. Samples were then dehydrated by a series of two minutes incubations in 70%, 85% and 100% ethanol, air dried and mounted using vectashield, containing DAPI (1 μg/ml). Intact and aberrant loci defined as red/green (yellow) fusion signals or split signals with a distance of more than one signal diameter, respectively, were evaluated and counted with a Leica DMI 4000 microscope. Statistical analysis of triplicate experiments was done using the Prism5 software (GraphPad, USA).

## ACCESSION NUMBERS

PAR-CLIP, RNA-seq and ChIP-seq libraries have been deposited in the Sequence Read Archive (SRA) under accession number SRP066889.

## SUPPLEMENTAL INFORMATION

## ACKNOWLEDGMENTS

We thank Manuel Stucki (University of Zürich) for his assistance in the evaluation of the 53BP1 foci, A. Kanitz, A. Rzepiela and R. Gumienny for contributing scripts, and S. Piscuoglio, M. Kovac, K. Heinimann and lab members (Department of Biomedicine, University of Basel), and Zavolan lab members for helpful discussions.

This works was supported by the grant KFS-3508-08-2014 from the Swiss Cancer League.

---

**Supplementary Figure 1.**
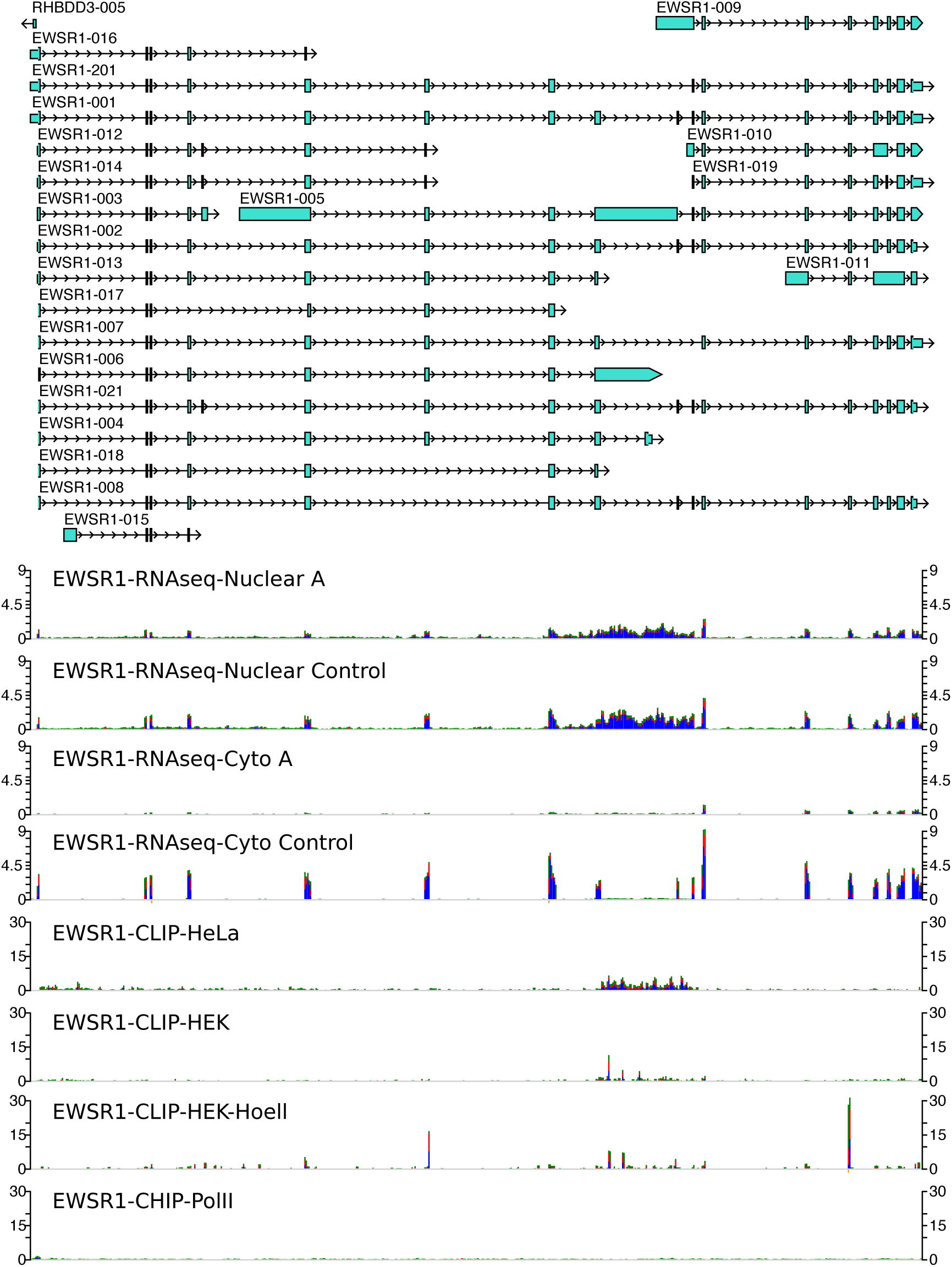
Screenshot of EWSR1 locus, showing annotated transcripts from the ENSEMBL database (top of the panel) and tracks illustrating the density of reads along the locus in the mRNA-seq samples prepared from the nucleus and cytoplasm of control siRNA and EWSR1 siRNA A-treated cells, PAR-CLIP samples prepared in our laboratory from HeLa and HEK cells, a sample from the study of Hoell et al. of HEK 293 cells over-expressing the EWSR1 protein (project accession SRA025082, sample accession SRS117978) and the PolII CHIP of EWSR1 from HeLa cells.

**Supplementary Figure 2.**
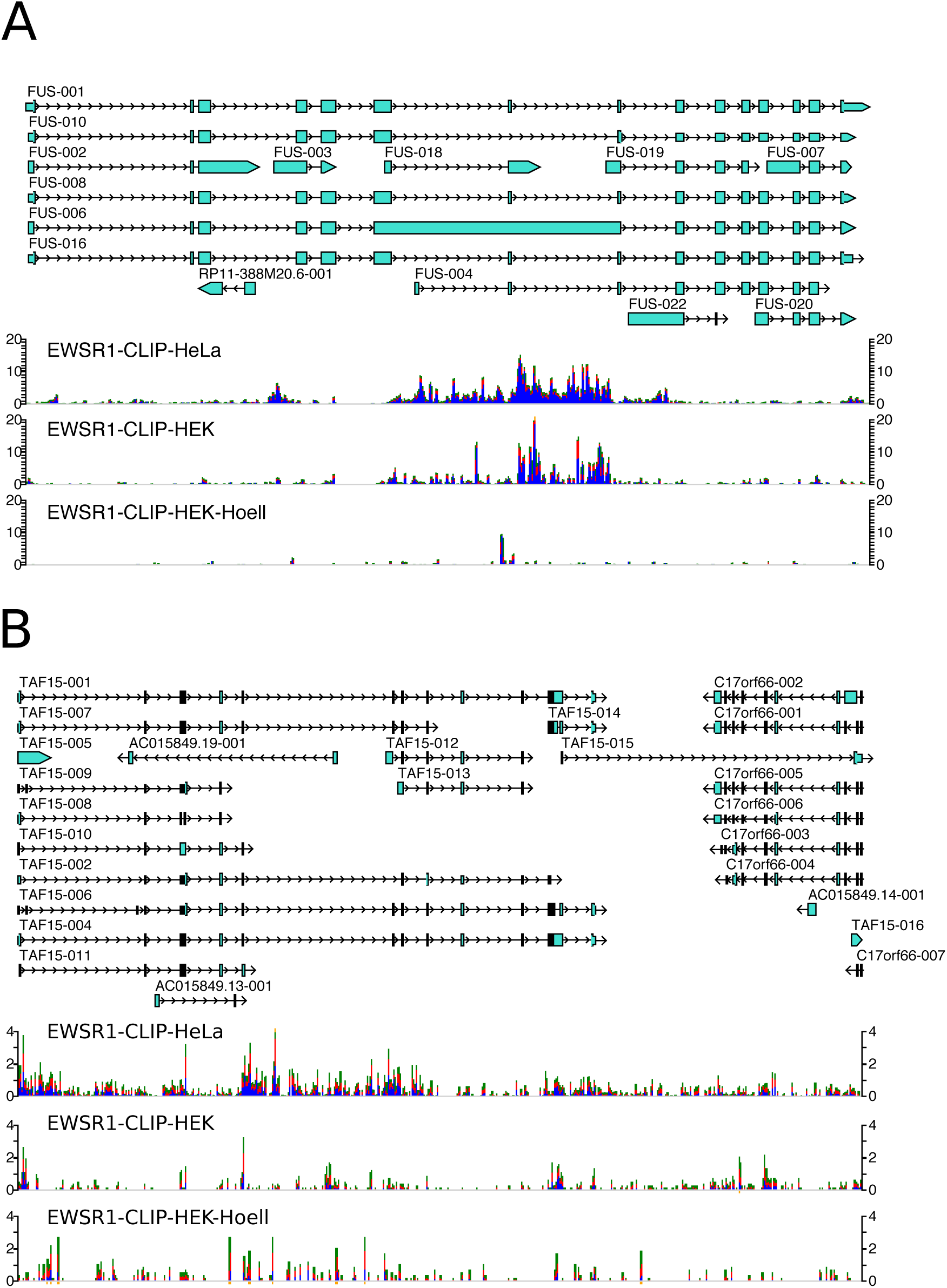
Screenshot of FUS and TAF15 loci, with a set of annotated transcripts from the ENSEMBL database (top of the panel) and the tracks illustrating data from different experiments, as generated by the clipz genome browser. The tracks show the density of reads along the locus in the, PAR-CLIP samples prepared in our laboratory from HeLa and HEK cells, a sample from the study of Hoell et al of HEK 293 cells over-expressing the EWSR1 protein.

**Supplementary Figure 3.**
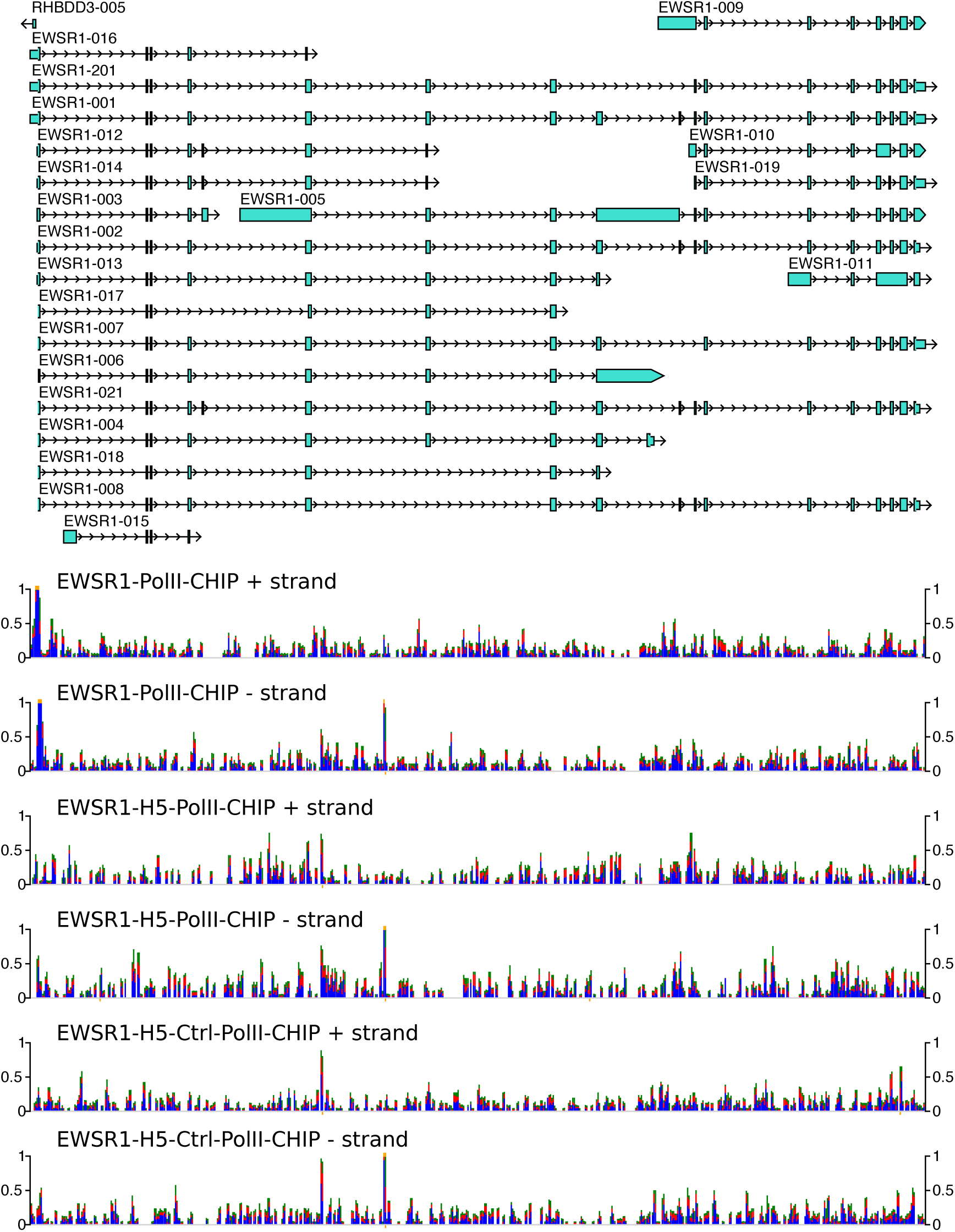
Density of reads in CHIP-seq experiments for RNAPolII (PolII) in untreated cells as well as its Ser2-phosphorylated form (H5-PolII) in cells treated with control or EWSR1-specific siRNAs.

